# Probing chemotaxis activity in *Escherichia coli* using fluorescent protein fusions

**DOI:** 10.1101/367110

**Authors:** Clémence Roggo, Jan Roelof van der Meer

**Affiliations:** Department of Fundamental Microbiology, University of Lausanne, 1015 Lausanne, Switzerland

## Abstract

Chemotaxis is based on ligand-receptor interactions that are transmitted via protein-protein interactions to the flagellar motors. Ligand-receptor interactions in chemotaxis can be deployed for the development of rapid biosensor assays, but there is no consensus as to what the best readout of such assays would have to be. Here we explore two potential fluorescent readouts of chemotactically active *Escherichia coli* cells. In the first, we probed interactions between the chemotaxis signaling proteins CheY and CheZ by fusing them individually with non-fluorescent parts of a ‘split’-Green Fluorescent Protein. Wild-type chemotactic cells but not mutants lacking the CheA kinase produced distinguishable fluorescence foci, two-thirds of which localize at the cell poles with the chemoreceptors and one-third at motor complexes. Cells expressing fusion proteins only were attracted to serine sources, demonstrating measurable functional interactions between CheY~P and CheZ. Fluorescent foci based on stable split-eGFP displayed small fluctuations in cells exposed to attractant or repellent, but those based on an unstable ASV-tagged eGFP showed a higher dynamic behaviour both in the foci intensity changes and the number of foci per cell. For the second readout, we expressed the pH-sensitive fluorophore pHluorin in the cyto- and periplasm of chemotactically active *E. coli*. Calibrations of pHluorin fluorescence as a function of pH demonstrated that cells accumulating near a chemo-attractant temporally increase cytoplasmic pH while decreasing periplasmic pH. Both readouts thus show promise as proxies for chemotaxis activity, but will have to be further optimized in order to deliver practical biosensor assays.

**IMPORTANCE:** Bacterial chemotaxis may be deployed for future biosensing purposes with the advantages of its chemoreceptor ligand-specificity and its minute-scale response time. On the downside, chemotaxis is ephemeral and more difficult to quantitatively read out than, e.g., reporter gene expression. It is thus important to investigate different alternative ways to interrogate chemotactic response of cells. Here we gauge the possibilities to measure dynamic response in the *Escherichia coli* chemotaxis pathway resulting from phosphorylated CheY-CheZ interactions by using (unstable) split-fluorescent proteins. We further test whether pH differences between cyto- and periplasm as a result of chemotactic activity can be measured with help of pH-sensitive fluorescent proteins. Our results show that both approaches conceptually function, but will need further improvement in terms of detection and assay types to be practical for biosensing.

## INTRODUCTION

Chemotaxis is the behaviour of cells to bias the direction of their motility in reaction to perceived chemical gradients (1, 2). Chemotaxis by bacteria is rapid (ms– to s–scale) and does not require *de novo* gene induction, since it is based on dynamic protein modifications and protein-protein interactions (3, 4). The rapidity of the chemotaxis response is potentially interesting for the development of alternative biosensor assays for chemical exposure testing. One current popular biosensor method relies on living bacterial cells equipped with synthetic gene circuits, which enable *de novo* expression of an easily-measurable reporter protein upon contact to target chemicals (5, 6). Gene expression, however, takes on average some 30 min to a few hours to show sufficient signal in the assay (7), which might be significantly shortened by interrogation of chemotactic response. Complicating for the deployment of chemotaxis as biosensor readout is that it is rather difficult to calibrate and quantify the response of chemotactically active motile cells (8). Firstly, it is tricky to produce a robust assay with stable chemical gradients that are a prerequisite for a calibrated chemotaxis response (9). Secondly, the chemotactic reaction itself can be quantified in a variety of ways. Chemotaxis can be deduced from accumulating cells in chemical gradients, for example, in capillary assays (10, 11), microfabricated chambers (12), agarose plug sources (13), or microfluidic platforms (14-16). Alternatively, assays can be based on Föster resonance energy transfer (FRET) measurements of dynamic interactions between fluorescently-labeled protein partners in the chemotaxis signaling pathway (17, 18).

Chemotaxis signaling in *Escherichia coli* at the molecular level starts by ligand-binding at the methyl-accepting chemotaxis receptors (MCPs) and is transmitted to the flagellar motor (Fig. 1A) (2-4). The MCPs phosphorylate the associated kinase protein CheA, which on its turn phosphorylates the response regulator protein CheY. The phosphorylated form of CheY (CheY~P) interacts with the FliM proteins of the flagellar motor, leading to a change in the direction of the motor from counterclockwise (CCW) to clockwise (CW) rotation (19-22). Rotation of the flagellar motor is accomplished by ion influx through the cytoplasmic membrane as source of energy. Some marine bacteria, such as *Vibrio* sp., use sodium motive force and Na^+^-influx, but the majority of motile bacteria like *E. coli*, uses the proton gradient and H^+^-influx (19, 23). The protons cross the membrane through up to eight protein complexes called the ‘stators’, composed of the MotA and MotB subunits that transmit the energy to the motor (19, 21). Steady dephosphorylation of CheY~P by the phosphatase CheZ maintains a constant ratio of CheY~P/CheY, and a basal level of alternating CCW and CW rotations. Attractant binding to the MCPs inhibits CheA kinase activity, which temporally decreases the level of CheY~P (Fig. 1A). A lower CheY~P/CheY ratio on average leads to a decrease of CW rotations, less tumbling and more straight runs, moving the cells upwards in the attractant gradient (2, 4, 24). On the contrary, binding of a repellent activates CheA, leading to an increase of CheY~P/CheY ratio and an increase of tumble events. This increases the chance of the bacteria to move downward in the repellent gradient. A methylation feedback loop on the MCPs allows re-establishment of the basal CheY~P/CheY ratio when cells find themselves over prolonged times at constant chemo-attractant concentrations (25). Thus, CheY~P levels rapidly change in cells exposed to a sudden change of concentration of attractant or repellent, followed by a return to the initial state within 5–10 minutes due to the methylation of the receptors (26). The dynamic interaction between CheY~P and its phosphatase CheZ or FliM has been measured by FRET (17, 18), and by direct observation of single molecules to motor complexes (27), and can be calibrated as a function of the attractant concentration (17).

**FIG 1.**
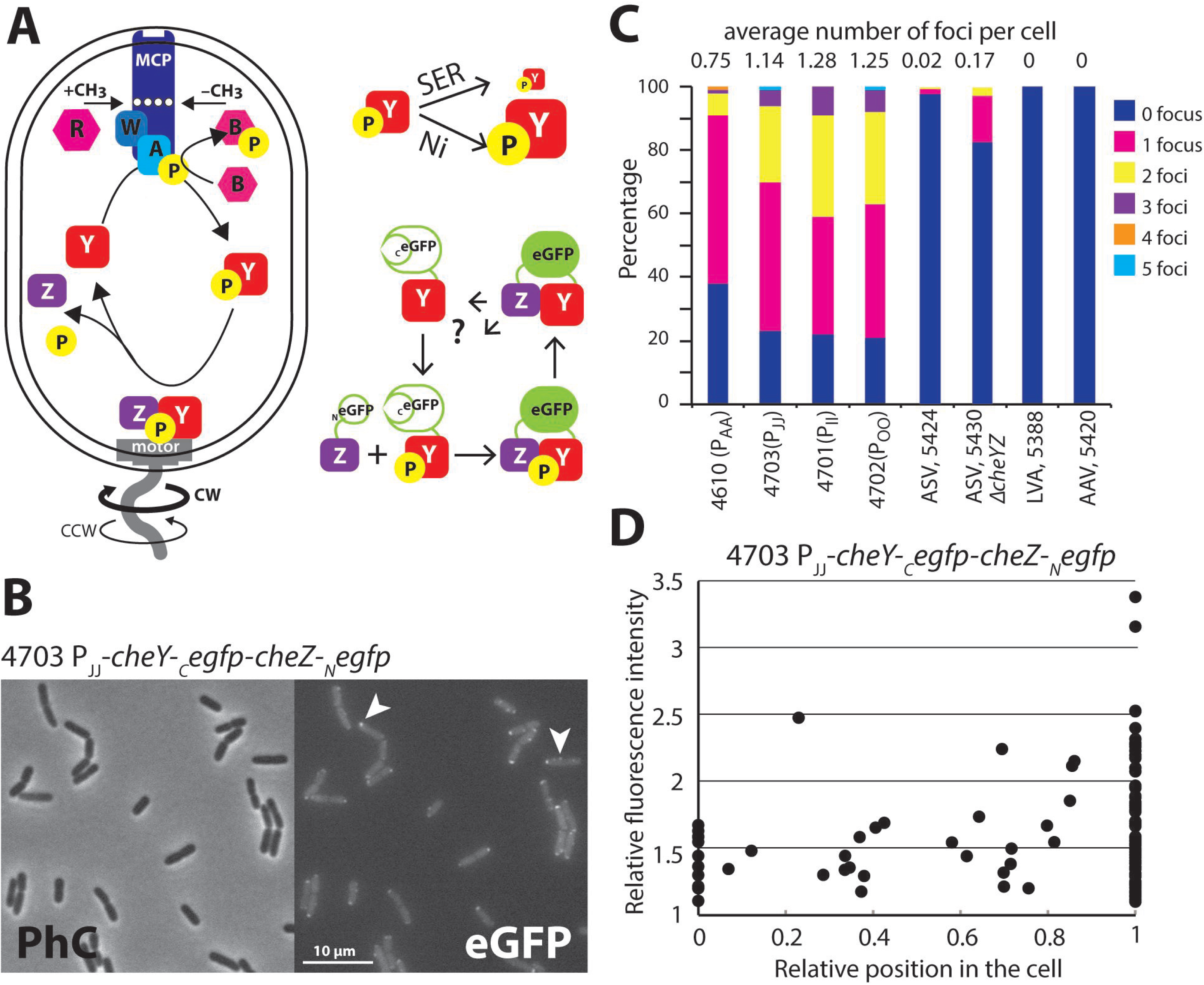
Bimolecular fluorescence complementation in chemotaxis. **(A)** Simplified scheme showing the major interacting partners in the *E. coli* chemotaxis signaling and the reconstitution of eGFP from interacting split versions of CheZ-_N_eGFP and CheY~P-_C_eGFP. Note how exposure to attractant (SER, serine) is expected to temporally reduce CheY~P pools, whereas exposure to repellent (Ni^2+^) is increasing those. **(B)** Visible formation of both polar as well as mid-cell eGFP foci (arrows) in *E. coli* strain 4703 expressing CheZ-_N_eGFP and CheY~P-_C_eGFP from the P_JJ_ promoter grown in absence of any chemo-attractant or -repellent. **(C)** Average number and distribution of eGFP foci numbers in a variety of *E. coli* strains differing in promoter driving CheZ-_N_eGFP and CheY~P-_C_eGFP expression, as well as carrying instability tags on eGFP. **(D)**. Superposition of eGFP foci positions and their relative intensity across n=100 cells of *E. coli* strain 4703.

In order to find potential alternative readouts for chemotactic behaviour of cells that might at some point enable development of biosensor assays, we decided to probe parts of the chemotactic signaling pathway and flagellar motor activity using fluorescence markers. In particular, we focused on quantifying CheY~P/CheZ interactions using bimolecular fluorescence complementation (BiFC), and secondly, we aimed to detect potential pH changes as a result of cellular motility using the pH-sensitive autofluorescent protein pHluorin (28). BiFC is based on the fusion of non-fluorescent parts of a fluorescent protein such as GFP to both proteins of interest that will reconstitute a functional fluorophore upon their interaction (29, 30). BiFC is frequently used for *in vivo* subcellular protein interaction localization in eukaryotes (31-33), but has also been deployed in bacteria (34-36). Although the generated split-GFP is stable, a few studies have shown a fast generation of fluorescence upon protein interactions in neurons (32, 37) and also signal decay (33, 37). Our hypothesis, therefore, was that we might localize CheY~P/CheZ interactions in chemotactic cells from reconstituted split-GFP, which might change in number, position or intensity depending on cells being attracted or repelled. Therefore we fused N- and C-terminal split parts of stable and unstable eGFP (38), respectively, to CheZ and CheY of *E. coli* (Fig. 1A). Formation of reconstituted eGFP foci was measured in a variety of mutant strains, to test the specificity and localization of foci formation. Dynamic foci behaviour (intensity and localization) in individual cells was quantified in conditions where cells were stimulated with chemo-attractants or -repellents.

As a second aim, we exploited the properties of pHluorin, a pH-sensitive variant of GFP that allows non-invasive and reversible measurements of intracellular pH (28). pHluorin shows a bimodal excitation at 390 nm and at 470 nm, and a single pH-dependent and reversible emission at 508 nm (28). The ratio of emission at 390-nm excitation divided by that at 470-nm excitation is linearly proportional to the pH (28). The precision for pH differences is approximately 0.2 pH units, measured on the cytoplasm of *Salmonella* (39) and *E. coli* (40). It has been shown that experimental manipulation of the intracellular and extracellular pH can modify flagellar rotation (41). We therefore speculated that pH-changes as a result of flagellar motor rotation differences in chemotactic cells might be observable from pHluorin emission ratio changes. pHluorin was expressed in the cytoplasm or the periplasm of motile *E. coli* and its fluorescence emission ratio was quantified as a function of external pH. Microscope assays were established to measured pHluorin emission ratio changes in populations of cells under conditions of active chemoattraction.

## RESULTS

**Expression of split-eGFP fused with CheY and CheZ chemotaxis proteins**. The chemotaxis protein CheY was fused with the C-terminal part of eGFP (amino acid: 158-238), and its phosphatase CheZ was fused with the complementary N-terminal part (amino acid: 2-157). We hypothesized that the interaction between phosphorylated CheY (CheY~P) and CheZ would favor binding of the split parts, inducing proper folding and eGFP fluorescence emission (Fig. 1A). Both fusion proteins were expressed from a single plasmid-located operon in *E. coli* under control of a constitutive synthetic promoter. Several promoters were tested, with an approximate difference in “strength”: P_AA_ > P_JJ_ > P_II_ > P_OO_ (strains 4610, 4703, 4701 and 4702, respectively) (42). Clear fluorescence foci were detected in absence of any added attractant or repellent in *E. coli* cells expressing both fusion proteins (Fig. 1B), from each of the four different promoters (Fig. S1), but not in *E. coli* containing the empty plasmid vector (strain 4717, Fig. S1). Strains carrying plasmids in which frameshift mutations were introduced into either *cheY* (strain 4728) or *cheZ* (strain 4729), or both (strain 4743), causing premature stop codons, did not show any fluorescent foci either (Fig. S1). We thus concluded that the foci were the genuine result of CheY-_C_eGFP/CheZ-_N_eGFP interactions and not the result of spontaneous split-eGFP reconstitution and subsequent multimerization.

Global fluorescence intensities (i.e., averaged across the whole cell) of *E. coli* expressing *cheY-_C_egfp-cheZ-_N_egfp* were not significantly higher than in the control strains (e.g., 4717, 4728, 4729, 4743), except for *E. coli* strain 4610 expressing *cheY-_C_egfp-cheZ-_N_egfp* from the strongest promoter P_AA_ (Fig. S2). This strain also showed the highest level of foci fluorescence, in comparison to *E. coli* expressing *cheY-_C_egfp-cheZ-_N_egfp* from P_OO_, P_JJ_ or P_II_ (Fig. S2). The number of foci varied between 0–5 per cell (Fig. 1C). In particular *E. coli* strain 4610 (P_AA_) showed fewer foci than strains 4701–4703 (P_II_, P_OO_ and P_JJ_, respectively) and in most cells only a single (polar) focus was observed (Fig. 1C, Fig. S2). The numbers of foci per cell were very similar for the strains 4703 (P_JJ_), 4701 (P_II_) and 4702 (P_OO_, Fig. 1C). In all strains expressing both fusion proteins simultaneously, two-thirds of the foci localized at the cell poles, and one-third at various positions along the cell (Fig. 1D, Fig. S2). The intensity of the foci was variable with a maximum of 3.5 times higher than the cell background, suggesting different accumulation or turnover of the fusion proteins at those sites. Adding instability tags to the _C_eGFP-part fused to CheY largely resulted in complete disappearance of foci in the *E. coli* strains (Fig. 1C, strains 5388 LVA-tag, and 5420 AAV-tag), but with 2% of cells still showing visible fluorescent foci in *E. coli* 5424 (ASV-tag).

**Functionality of the split-eGFP fusion proteins**. In order to test whether the split-eGFP chemotaxis fusion proteins were functional, we cloned pCRO9 (P_JJ_*-cheY-_C_egfp-cheZ-_N_egfp*) into an *E.coli* Δ*cheYcheZ* deletion mutant (strain 5391, which is not chemotactic, Fig. 2A). In this *E. coli* background, the chemotaxis regulators will only be expressed as split-eGFP fusion protein. Cell accumulation after 30 minutes of *E. coli* 5395 (pCRO9 in Δ*cheYcheZ*) close to a solid source containing 100 μM serine was less steep and more diffuse than *E. coli* 4498 (MG1655 wild-type, Fig. 2A), but was otherwise comparable. This confirms that the fusion proteins are functional to control chemotaxis. The average number of foci in *E. coli* 5393 (pCRO9 in Δ*cheYcheZ*) in absence of chemotactic stimulation was statistically significantly higher than in strain 4703 (pCRO9 in MG1655; p=0.00059, pair-wise t-test; Fig. 2B). Also, the number of foci in *E. coli* 5430 (pCRO32 with ASV-destabilization tag on eGFP, in Δ*cheYcheZ*) was higher than in strain 5424 (pCRO32 in MG1655; Fig. 1C). The higher foci number is consistent with the idea that the split-eGFP fusion proteins in the Δ*cheYcheZ* mutant exclusively react among each other (creating foci), whereas in wild-type *E. coli* interactions of split-eGFP-versions with native CheY and CheZ protein result in non-fluorescent complexes.

**FIG 2.**
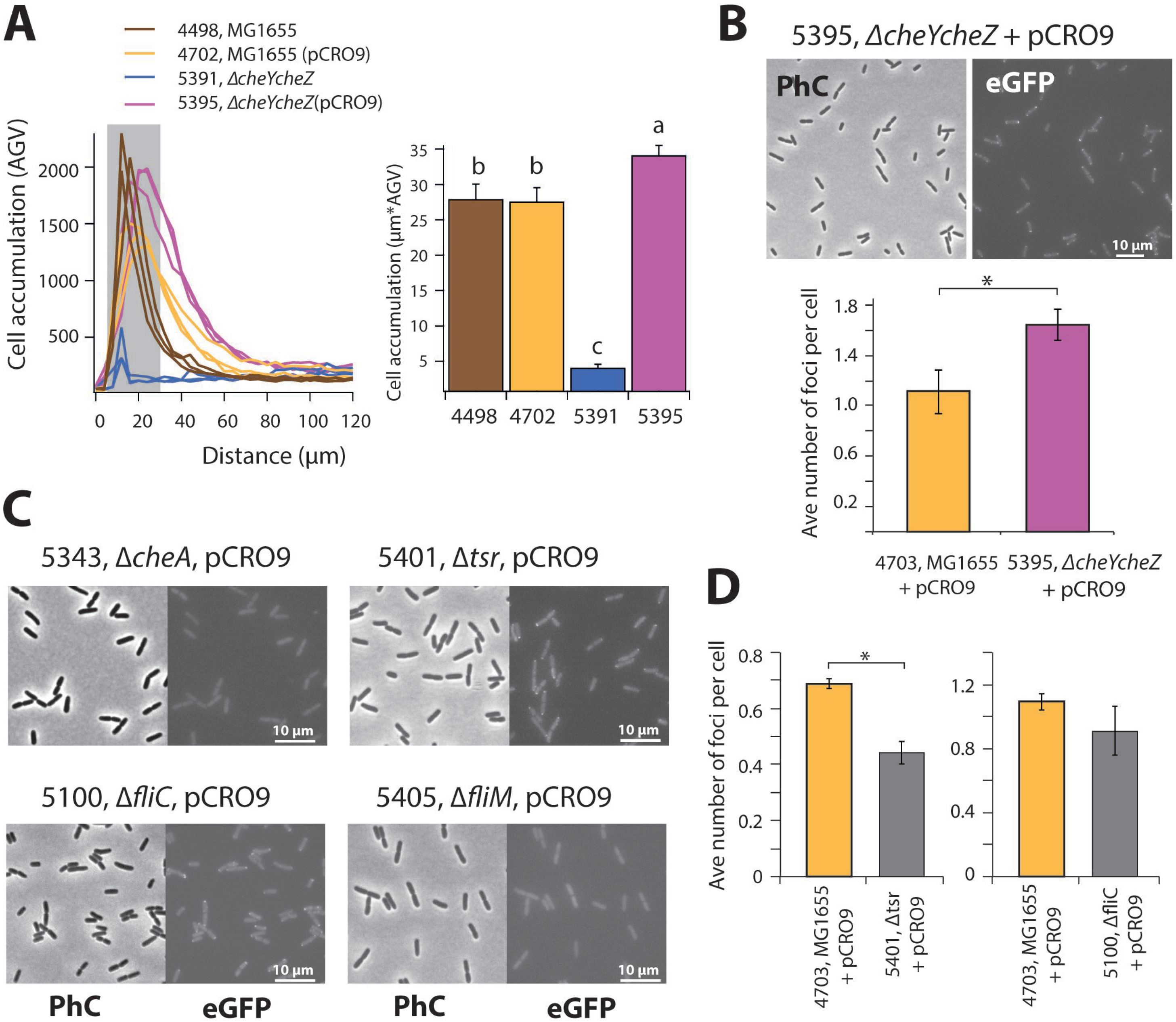
Functionality of CheZ-_N_eGFP and CheY~P-_C_eGFP split proteins. **(A)** Cell accumulation within 30 min to a 100-μM serine source of *E. coli* strains expressing *cheY-_C_egfp-cheZ-_N_egfp* (pCRO9) compared to MG1655 wild-type or MG1655 deleted of wild-type *cheY-cheZ*. Bar plots show cell accumulation within the first 30 μm closest to the source edge (in grey in the left panel). Letters indicate significance groups in ANOVA (p<0.05). **(B)** Increased average number of reconstituted eGFP foci in *E. coli* 5395 in absence of chromosomal *cheY-cheZ* compared to wild-type (*, p=0.0059). **(C)** Effect of deletions of *cheA, tsr, fliC* or *fliM* on appearance of reconstituted eGFP foci in *E. coli* carrying plasmid pCRO9 with *cheY-_C_egfp-cheZ-_N_egfp*. **(D)** Average number of eGFP foci per cell in Δ*fliC* or Δ*tsr* mutants compared to wild-type *E. coli*. *, p=0.023, pair-wise two-tailed t-test.

**Split-eGFP expression in chemotaxis deletion mutants**. In order to confirm that the foci detected were a consequence of interaction between CheY~P and CheZ, we introduced the P_JJ_-*cheY*-*_C_egfp-cheZ-_N_egfp* (pCRO9) construct into an *E. coli* mutant lacking the gene encoding the CheA kinase. Deletion of CheA abolishes phosphorylation of CheY and of the CheY-_C_eGFP fusion protein, which should cancel interactions with the CheZ phosphatase. *E. coli* Δ*cheA* (pCRO9) cells indeed did not show any foci (Fig. 2C), which is in agreement with our hypothesis and confirms that the observed foci in e.g., *E. coli* strain 4703 or 5395 must be the result of physical interaction between phosphorylated CheY-_C_eGFP and CheZ-_N_eGFP.

Since CheY~P–CheZ-split-eGFP fluorescence mostly appeared in clearly localized foci in the cell (Fig. 2B), this suggested they were the result of an additional interaction of either CheY~P or CheZ with either chemoreceptors or motors, which have precise localization (43). CheY/CheY~P is known to interact both with the chemoreceptors and the flagellar motors (44). In an *E. coli* Δ*tsr* mutant carrying pCRO9, which lacks the major chemoreceptor for serine, fluorescent foci were still visible (Fig. 2C), but the average number of foci per cell was statistically significantly reduced compared to wild-type *E. coli* strain 4703 with pCRO9 (p=0.023, pair-wise t-test; Fig. 2D). This suggested that split-eGFP foci at least partially arise at the chemoreceptor positions. In an *E. coli* without *fliM* split-eGFP foci could no longer be detected at all (Fig. 2C). The *fliM* gene encodes the motor protein with which CheY~P interacts to invert flagellar rotation, but absence of FliM also prevents export of the anti-sigma factor FlgM (45). This impedes expression of the downstream flagellar genes, including the chemoreceptors, native *cheY/Z* and *cheA/W* (45). This indicated that no spurious foci form outside chemoreceptors and flagellar motors. In absence of flagella (*fliC* deletion mutant) the average number of foci per cell was only slightly lower than in wild-type motile *E. coli*, but this difference was not statistically significant (Fig. 2C, D).

To provide additional evidence that CheY~P-_C_eGFP and CheZ-_N_eGFP also interact at the flagellar motors, we constructed an *E. coli* MG1655 derivative strain co-expressing the CheY-CheZ split-eGFP fusion proteins and a FliM-mCherry fusion protein (strain 5921, Table 1). Fluorescent foci of both eGFP and mCherry were visible in individual cells, although in general the eGFP foci were a bit more ‘crisp’ due to lower background than in case of the mCherry foci (Fig. 3A). Automated segmentation of cells and foci indicated a larger average number of mCherry than eGFP foci per cell (Fig. 3B, C). eGFP foci, as before, were most frequently located at the cell poles, although consistently about one-third of foci was found closer to the mid-cell (Fig. 3B). In contrast, two-thirds of all FliM-mCherry foci were found along the mid-cell (Fig. 3B), suggesting an on average larger number of spatially distinct flagellar motor than receptor complexes (at the resolution of regular epifluorescence microscopy). At focal centre distances of less than 120 or 180 nm, which is the resolution of epifluorescence microscopy, there was 22% or 37%, respectively, of eGFP foci overlapping with FliM-mCherry foci, one-third of which were localized in mid-cell (Fig. 3C). This strongly suggests that part of the CheY~P-_C_eGFP-CheZ-_N_eGFP complexes assemble at the flagellar motors.

**FIG 3.**
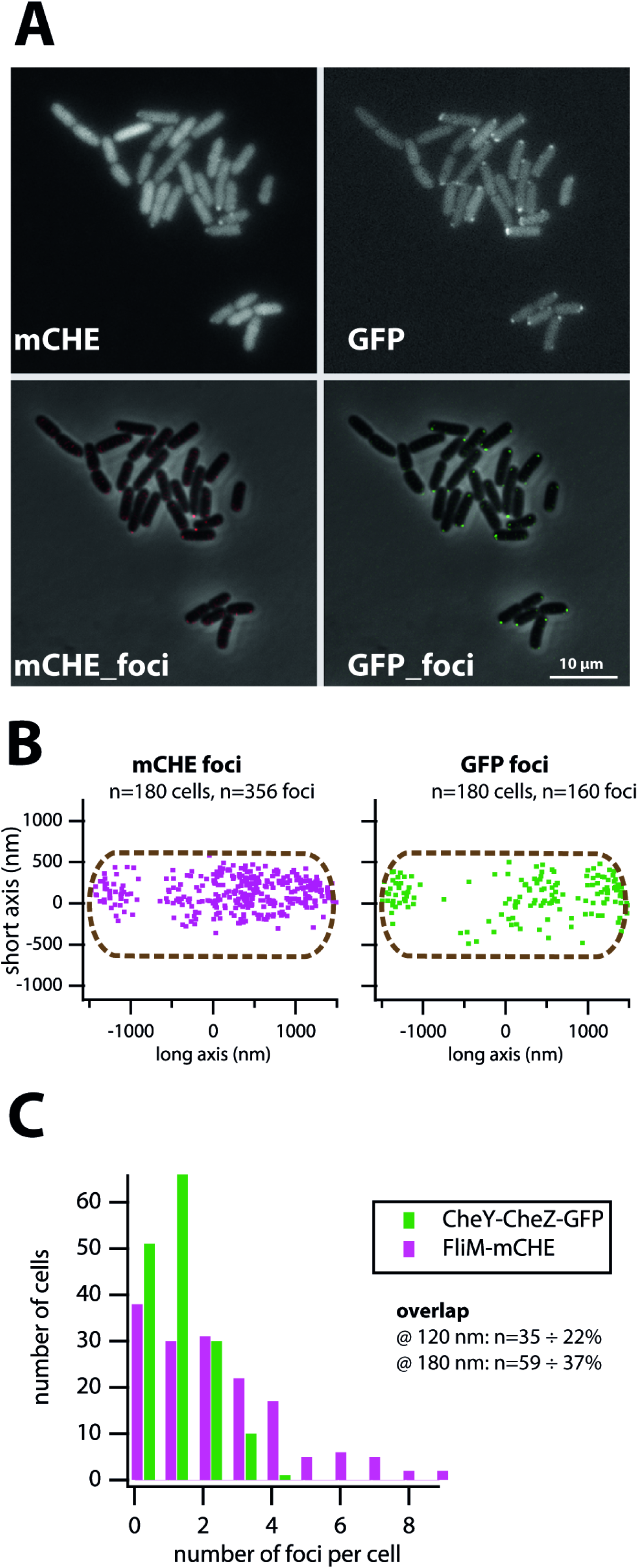
Colocalization of reconstituted CheY-CheZ-eGFP foci with FliM-mCherry. **(A)** Representative images of *E. coli* 5921 expressing both FliM-mCherry and *cheY-_C_egfp-cheZ-_N_egfp* from P_JJ_ (on pCRO9, top series), and after foci detection (local background subtraction by SuperSegger analysis, lower series). **(B)** Superposition of FliM-mCherry and Split-eGFP foci positions among 180 cells, plotted along a normalized *E. coli* cell. **(C)** Distribution of number of FliM-mCherry and split-eGFP foci per cell and the proportion of overlapping foci at less than 2 or 3 pixel resolution (120 or 180 nm).

**Table 1.**
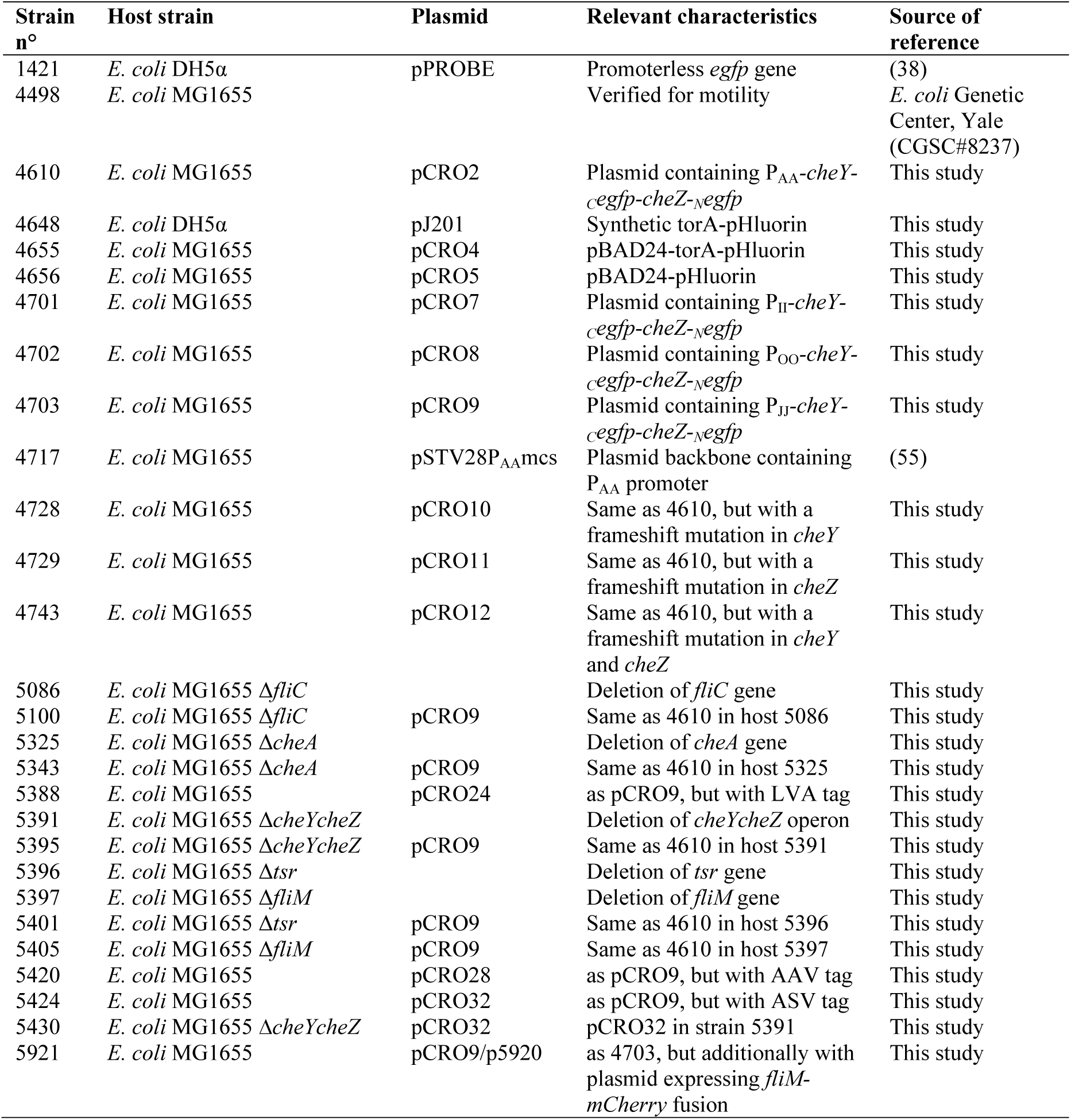
Strain list

**Dynamic response of CheY-CheZ split-eGFP foci**. In order to measure potential dynamic changes in CheY~P-_C_eGFP-CheZ-_N_eGFP foci as a result of chemotactic response, we immobilized *E. coli* Δ*cheYcheZ* (pCRO9) cells expressing both fusion proteins on the bottom of poly-L-lysine coated wells on top of a microscopy slide immersed in motility buffer. Foci fluorescence was recorded by microscopy imaging every minute, while focusing on cells attached to the slide, before and after addition of ligand (100 μM Ser or Ni^2+^). Fluorescence response curves of individual foci were variable and mostly decreased over time, as a consequence of photobleaching (Fig. 4A). In comparison to the average fitted slope of foci fluorescence decay (Eq. 1) in *E. coli* cells remaining in motility buffer, the proportion of foci with a significantly slower decay was higher in cells exposed to 100 μM Ni^2+^ (p=0.0072, Kruskal-Wallis test, Fig. 4B). Slower decrease than expected from photobleaching might be the result of faster renewed CheY-phosphorylation and new formation of split-eGFP in repelled cells. In contrast, the proportion of foci with faster fluorescence decay than expected from motility buffer was not significantly different in cells to which 100 μM serine was added. Conversely, neither Ni^2+^ or serine addition resulted in different proportions of response curves significantly higher than the average in motility buffer (Fig. 4B).

**FIG 4.**
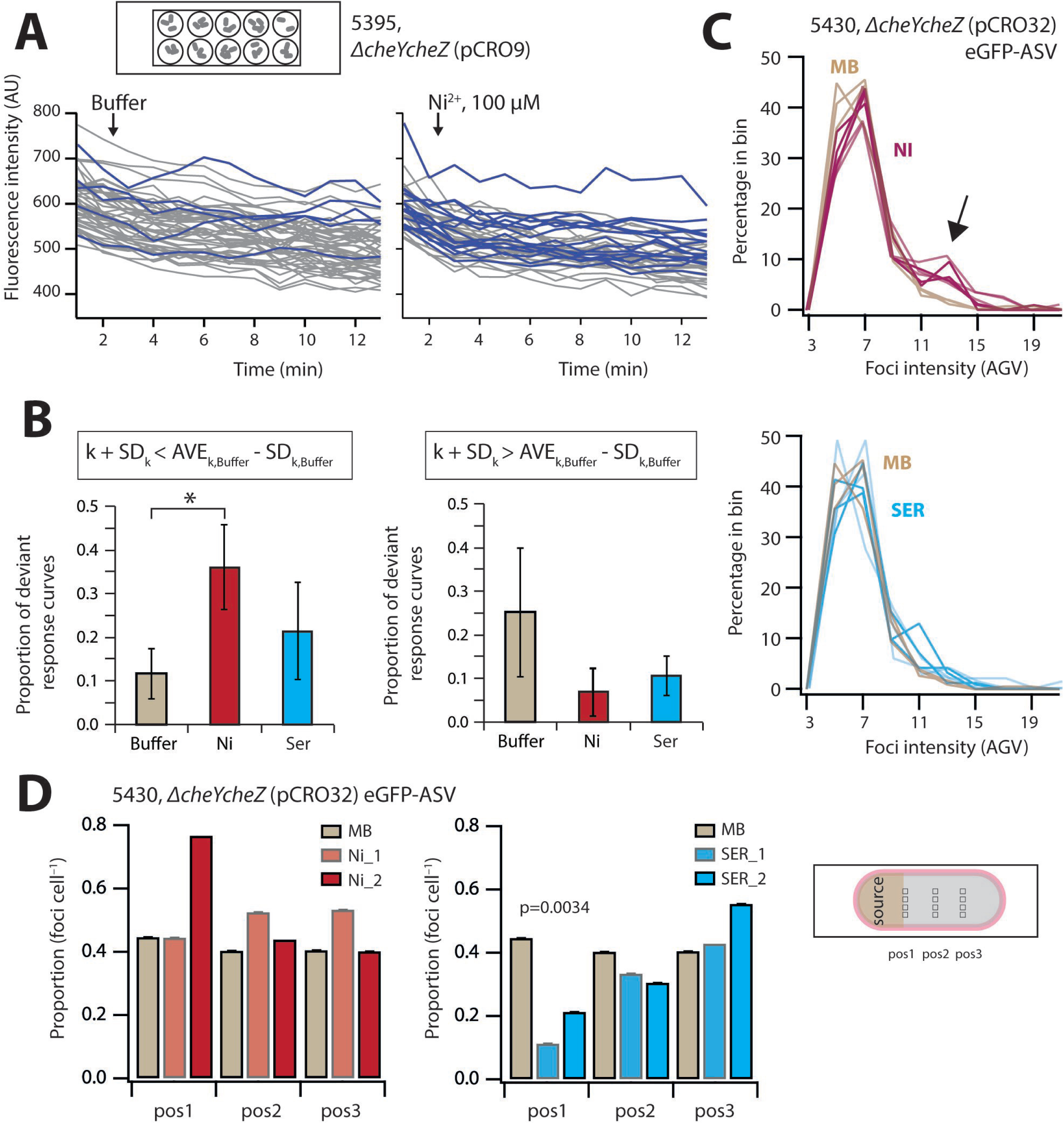
Split eGFP foci dynamism after ligand exposure. **(A)** Foci fluorescence of surface-immobilized *E. coli* Δ*cheY*Δ*cheZ* (pCRO9) cells in wells with motility buffer or exposed to 100 μΜ NiCl_2_ after time 2 min. Blue curves indicate foci with less fluorescence decay than expected from the mean slope of all decay curves. **(B)** Proportion of deviant foci response curves compared to the mean slope of all foci decay curves in motility buffer ± one SD (*, p=0.0072). **(C)** Foci intensity distributions among surface-spread *E. coli* 5430 cells expressing unstable split-eGFP at three distances on a solid source with either 100 μΜ NiCl_2_, 100 μΜ serine (SER), or motility buffer (MB) alone, each in two replicates. Note that distances are not differentiated here. Note the shift to brighter foci in Ni^2+^-exposed cells with values of 11-17 AGV (arrow). **(D)** Average proportion of foci per cell for the experiments in (C) as a function of position. Decreasing foci numbers as a function of distance (pos3 to pos1) for serine-exposed cells is statistically significant (LINEST slope test, p=0.0034), whereas those in cells exposed to Ni^2+^ or in MB do not differ significantly from the Null hypothesis (Slope = 0).

Comparison of fluorescence intensity distributions of stable split-eGFP foci in immobilized *E. coli* Δ*cheY*Δ*cheZ* (pCRO9) cells in the wells over time (Fig. S3), showed a slight increase towards brighter foci in case of cells exposed to Ni^2+^ compared to motility buffer, although this was not statistically significant (Fig. S4). For cells exposed to serine, the proportion of weaker foci tended to increase (Fig. S4). In case of immobilized *E. coli* Δ*cheY*Δ*cheZ* (pCRO32) producing unstable split–eGFP-ASV, more brighter foci appeared over time after exposure to Ni^2+^, whereas no consistent changes in foci distribution appeared after exposure to serine (Fig. S4).

In the next experiment, instead of being attached to wells filled with motility buffer, *E. coli* Δ*cheY*Δ*cheZ* (pCRO32) cells were deposited on agarose surfaces at different distances from either a source with 100 μM serine, 100 μM Ni^2+^, or in motility buffer only. Exposure to Ni^2+^ led to a net increase in the proportion of brighter foci compared to motility buffer, whereas exposure to serine did not measurably change foci brightness’ distribution (Fig. 4C). In contrast, the overall number of foci per cell decreased in case of cells upstream in the serine gradient (Fig. 4D, p=0.0034 compared to Null hypothesis of no decrease).

Collectively, these results indicate that the changes in dynamic CheY~P–CheZ interactions as a result of exposure to attractant or repellent can be captured to some extent by split-eGFP foci formation, although the response of individual cells is highly heterogeneous and the difference between conditions is small.

**pHluorin captures pH-changes in chemotactic cells**. Our second aim was to determine possible pH changes in the peri- and cytoplasm of *E. coli* during or as a consequence of altered swimming behaviour in an attractant gradient, for which we deployed the pH-sensitive fluorophore pHluorin. pHluorin expressed from the L-arabinose inducible *araC* promoter in wild-type motile *E. coli* MG1655 at a concentration of 1 g l^−1^ arabinose showed homogenous fluorescence distribution in the cytoplasm (Fig. 4A). Fluorescence intensity of TorA-pHluorin was highest at 10 g l^−1^ arabinose, with cells showing a “ring-like” appearance, indicating that the protein is efficiently exported into the periplasm (Fig. 4A, Fig. S5). The average ratio of 525-nm emission at 386-nm divided by that at 470-nm excitation of pHluorin as well as TorA-pHluorin increased linearly between pH 6 and 8 in individual *E. coli* cells observed on agarose patches containing 20 mM sodium benzoate at pHs ranging between 6 and 8 (Fig. 4B). The resulting calibration curve was then used to deduce the intracellular and periplasmic pH of *E. coli* cells during chemoattraction.

Because of the difficulty to measure pHluorin fluorescence ratios in motile cells, we adapted the agarose plug assay to maximize the number of chemotactically active cells in microscopy settings (Fig. 4C). Washed suspensions of *E. coli* expressing pHluorin or TorA-pHluorin inserted in a very shallow (0.17 mm) microscope chamber filled with a round and flat solidified agarose source containing 0.1 mM serine clearly accumulated close to the source 15–20 min after the start of the incubation (Fig. 4C). No significant cell accumulation was observed with an agarose source that did not contain any attractant (Fig. 4C). *E. coli* cell populations expressing pHluorin (in the cytoplasm) showed an increase of the pHluorin emission ratio close to the serine source followed by a general decrease, compared to a constant signal decrease in absence of any attractant (Fig. 4D). In contrast, *E. coli* cell populations expressing TorA-pHluorin showed a steep decrease of the emission ratio close to the source followed by a slow increase, compared to a constant signal decrease in the control (Fig. 4D). In both instances, cells furthest away from the source showed equal emission ratios for attractant or no attractant. These measurements would imply an increase of the pH in the cytoplasm and decrease in the periplasm of motile cells attracted in a chemical gradient.

## DISCUSSION

The goal of this work was to develop different readouts of the chemotaxis pathway of *E. coli*, which might be used as proxies for chemotactically active cell behaviour, and which might eventually be exploited for biosensing. Although other approaches have been taken, we focused here on two types of reactions. In the first, we followed interactions between CheY~P and CheZ as a proxy of ligand binding to the chemoreceptors in *E. coli* using BiFC with split-eGFPs. In the second approach, we studied potential dynamic pH-changes in chemotactically active cells using the pH-sensitive fluorophore pHluorin.

Our results demonstrated that CheY-_C_eGFP and CheZ-_N_eGFP fusion proteins were functionally complementing chemotaxis in an *E. coli* Δ*cheYcheZ* deletion mutant background (Fig. 2). Chemo-attraction of *E. coli* expressing CheY-_C_eGFP and CheZ-_N_eGFP was slightly less steep than that of *E. coli* MG1655 WT, both in presence or absence of native CheY/CheZ (Fig. 2). This suggests that the fusion proteins not measurably interfere with the native proteins or impair chemotaxis. Clear eGFP fluorescent foci were present in *E. coli* expressing simultaneously the CheY-CeGFP and CheZ-NeGFP fusion proteins, which were both localized at the cell poles as well as at different positions along the cell length (Fig. 1 and 2). Deletion of the gene for the CheA kinase that phosphorylates CheY abolished formation of any foci (Fig. 2), which demonstrated that the foci are the result of specific interactions between CheY~P and CheZ, since non-phosphorylated CheY and CheZ do not interact. We found that eGFP foci colocalize both with the chemoreceptors and with the flagellar motors (Fig. 3), both of which are known sites for CheY~P interaction (44). Expression of CheY-CeGFP and CheZ-_N_eGFP in a Δ*tsr E. coli* mutant background showed a decrease in the average number of foci per cell, but did not abolish foci formation altogether. This suggests that Tsr partially stabilizes CheY~P–CheZ interaction (Fig. 2D). No foci were detected in *E. coli* Δ*fliM*, but this is not (only) the result of direct absence of motor proteins stabilizing the interaction to CheY-_C_eGFP. In absence of FliM the anti-sigma factor FlgM is not transported from the cytoplasm, which blocks transcription of many chemotaxis genes, among others, of *cheY/Z* and *cheA* itself (45). Mathematical modeling has demonstrated that the polar localization of CheZ is crucial to obtain a uniform concentration of CheY~P in the cell in order to interact similarly with all peritrichous flagella (46). CheZ and CheY~P are thus supposed to interact at the receptor near the cell pole (47, 48). However, co-labeling with a FliM-mCherry fusion indicated that around one-third of split-eGFP fusions colocalize with the motor complexes (Fig. 3). Hence, we conclude that the CheY~P/CheZ interactions can take place both at the motor and at the receptor. The latter is consistent with theory because CheY and CheZ are localized at the receptor, as shown by Sourjik and Berg (44), but CheY~P, once phosphorylated by CheA, has to diffuse to the motor to induce inversion of flagellar rotation (27) and may therefore further interact with CheZ at the motor.

Although the levels of CheY~P in the cell are dynamic, since dependent on the binding of attractants or repellents to the receptors, and further on CheZ constantly dephosphorylating CheY~P, the CheY~P–CheZ–reconstituted–eGFP foci were rather stable. The cause for this is most likely the stability of the reconstituted eGFP itself. By photobleaching cells in motility buffer in comparison to motility buffer with addition of nickel ions (a strong repellent), we could see a trend that eGFP photodecay was counteracted by renewed foci formation (Fig. 4A, B; Fig. 6A). In cells exposed to serine we did not detect any significant changes of foci intensity compared to cells in motility buffer only, but this might be the result of the stability of split-eGFP-CheY-CheZ foci, which do not dissociate faster upon addition of attractant. When looking at the distribution of foci intensities among individual surface-attached cells in the wells, there was again a trend that Ni-exposed cells accumulated brighter foci over time (Fig. S4). The effects were more clear when using *E. coli* expressing the split-eGFP appended with the ASV-destabilization tag (strain 5430, Table 1) (49). In this case, foci are more rapidly degraded and the appearance of new brighter foci as in Ni^2+^-exposed cells can be more easily distinguished (Fig. S4). This was further confirmed by placing *E. coli* strain 5430 cells on agarose surfaces with gradients in Ni^2+^- or serine-concentration, which led to brighter foci appearing in Ni^2+^- exposed cells and an overall decrease of the number of foci in cells exposed to serine (Fig. 4C, D). These results would be in agreement with what one would conjecture from the expected changes in CheY~P levels (Fig. 6A). Both the foci brightness and the number of foci per cell thus reflect temporal chemotaxis pathway activation through modification of the CheY~P levels by the binding of attractant or repellent to the receptors: attractants inducing a decrease of the concentration of CheY~P and repellents leading to an increase (Fig. 6A).

Most publications deploying BiFC indeed indicate split-fluorescent proteins to be stable and irreversible (31), with some exceptions in eukaryotic cells (33, 37). Stability of the reconstituted eGFP thus prevents capturing much of the dynamic nature of CheY~P/CheZ interactions, which was improved by deployment of the ASV-tagged eGFP. However, the downside of the ASV-tagged split-eGFP is that its turnover is higher, and fluorescent foci on average become much weaker and more difficult to detect. Fluorescent foci detection is optimal on surface immobilized cells, but the immobilization itself might hinder flagella rotation and diminish CheY~P/CheZ interactions at the motors. A further problem in the current assay configuration is the addition of ligand to stimulate chemotaxis. It is difficult to estimate the time delay for the reaction of the cells after adding the ligand into the solution in case of the bacteria attached to the bottom of the wells (as in the configuration of Fig. 4A, B). Likewise, in that configuration it is difficult to estimate the duration of the transient reaction of the cells, until they adapt to the new ligand concentration as a result of methylation of the chemoreceptors. In case of the source assay as in Fig. 4C and D, a ligand gradient is formed along the agarose, and individual cells will react during a longer time because they remain within a concentration gradient. Other split-GFP variants may help to improve the system further. For example, a tripartite GFP was described recently which shows faster association and minimized non-specific protein aggregation (50). The use of dimerization-dependent fluorescent proteins (ddFP) could also be an interesting alternative. It consists of two weakly or non-fluorescent monomers of fluorescent protein that become fluorescent upon interactions (51, 52). The advantage is the reversibility of the system, which can react in a timescale of seconds (52). It is not known, however, if the ddFP monomers would affect the functionality of CheY and CheZ.

Our second approach consisted of measuring temporal pH differences as a result of chemotaxis. Flagellar rotation in *E. coli* is powered by a proton flux through the cytoplasmic membrane (41). We hypothesized that the addition of attractant might lead to a temporary decrease of the pH in the cytoplasm as a result of proton influx and pH increase in the periplasm as a result of temporary proton depletion (Fig. 6B). To measure this, we expressed the pH-sensitive pHluorin fluorescent protein in the cytoplasm and in the periplasm, and exposed *E. coli* cells for 10–15 min to a gradient diffusing from a solid agarose source containing 100 μΜ serine. The difficulty, however, was to obtain sufficient imaging quality of both accumulating cells and their fluorescence at two excitation wavelengths. Because of the time needed for filter change we could not image individual motile cells at high magnification (e.g., 600−1000×). Instead, we relied on lower magnifications, which are less sensitive to cell blurring (200×). Quite robust values of pHluorin and TorA-pHluorin emission ratios were obtained from accumulating *E. coli* cells as a function of distance to the solid source (Fig. 5C). Since cells are within a serine gradient, they will react during a relatively long time (10–25 min), enabling optimal measurements. Interestingly, the pHluorin emission ratio increased in cells close to the serine source compared to an empty source, whereas the TorA-pHluorin emission ratio decreased closed to the serine source (Fig. 5D). The inverse response in the cyto- versus periplasm, suggests that the pH of the cytoplasm increases whereas that of the periplasm decreases in chemotactically-active cells (Fig. 6B). The fact that both signals were opposite indicates that they were not an artifact simply due to the cell accumulation close to the source. Calculation of the cytoplasmic and periplasmic pHs from the pH–calibrated pHluorin fluorescence ratios (Fig. 5B) indicated that, in absence of attractant, the equivalent cytoplasmic pH corresponds to 7.8±0.16, which is 0.3 pH-unit higher than measured elsewhere (40, 53), and that of the periplasm to 7.3±0.12 (Fig. 6B). A lower pH in the periplasm is in general agreement with a net outside proton gradient across the cytoplasmic membrane in actively respiring cells (53). In contrast, in presence of an attractant, reactive cells showed an increase of 0.3 pH units in the cytoplasm (from pH 7.8 to 8.1) and a decrease of 0.2 units in the periplasm (from pH 7.3 to 7.1) (Fig. 6B).

**FIG 5.**
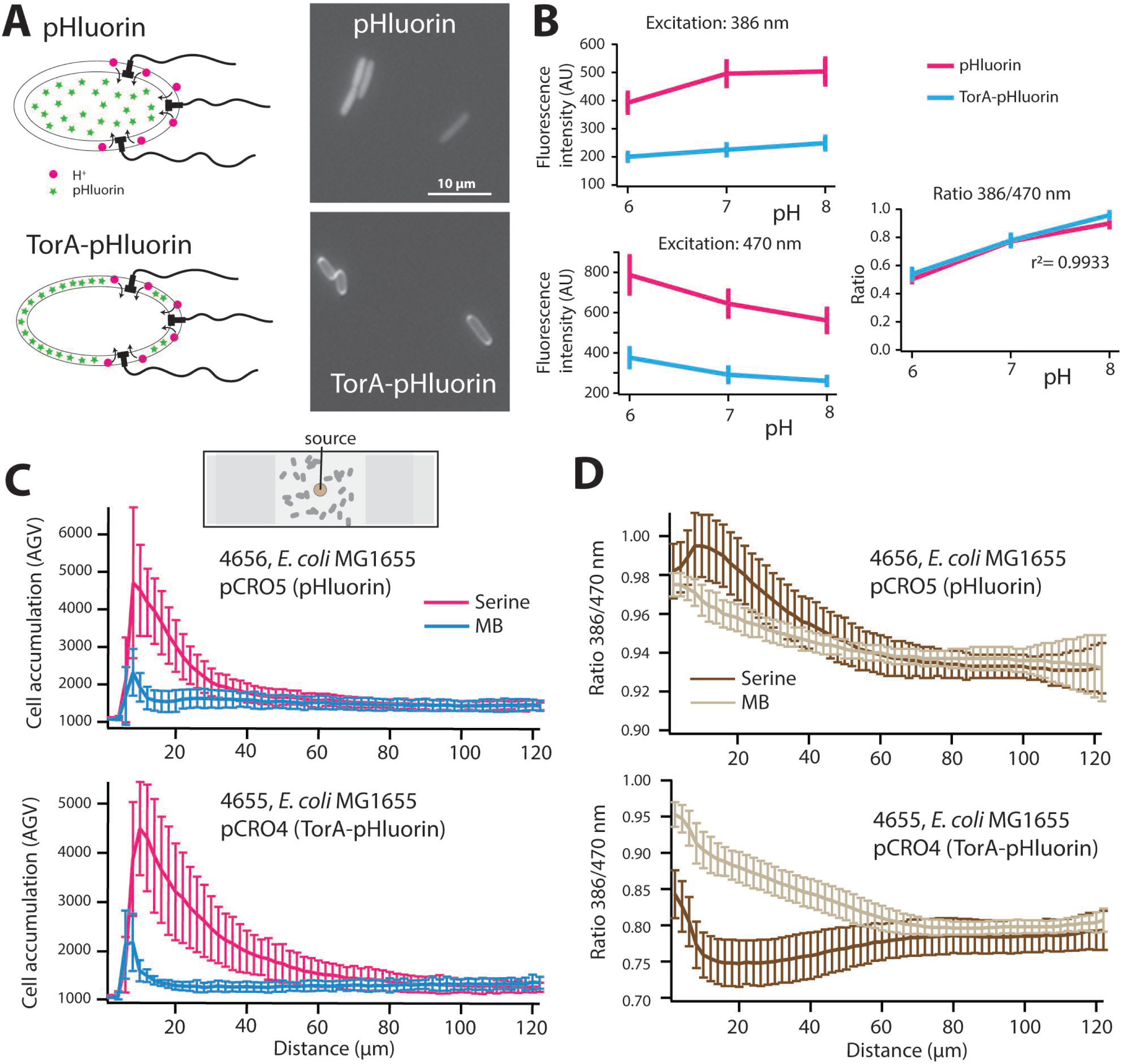
pH changes in chemotactically-active *E. coli* cells. **(A)** Expression of pHluorin or TorA-pHluorin in the *E. coli* cyto- and periplasm, respectively. **(B)** pHluorin excitation and emission ratios as a function of external pH. **(C)** Cell accumulation (top) and pHluorin emission ratios in *E. coli* 4656 (pCRO5) after 20 min as a function of distance to a 100−μM serine source (red data points) or a source with motility buffer (MB), imaged at 200× magnification in the set-up as schematically outlined. (D), as (C) but for *E. coli* strain 4655 (pCRO4), expressing TorA-pHluorin. Bars denote calculated standard deviations from the mean at that position in four biological replicates imaged on both sides of the agarose source.

These results are not in immediate agreement with our initial hypothesis that increased chemotaxis activity would increase proton flux through the stator of the motor, and, correspondingly, would decrease cytoplasmic pH. Instead, our results suggests that chemotactically-active cells increase proton efflux from the cytoplasm to the periplasm, perhaps in order to compensate and sustain the high proton influx through the flagellar motor. The increased proton efflux may find its origin in temporarily higher respiration rates (53). The net increase of pH gradient between the periplasm and the cytoplasm would on its turn facilitate the proton requirements by the flagellar motor for faster or continued rotation.

In conclusion, our study showed how autofluorescent proteins may be used to interrogate the chemotaxis pathway in motile *E. coli* in gradients of attractant or repellent. BiFC with unstable split-eGFP parts fused to CheY and CheZ can to some extent reveal the dynamic behaviour of CheY~P/CheZ foci (Fig. 6A), whereas pHluorin expressed in the cyto- and periplasm can measure dynamic pH changes in chemotactically attracted cells (Fig. 6B). Both methods may be further optimized and calibrated to allow quantitative chemotaxis readout.

**FIG 6.**
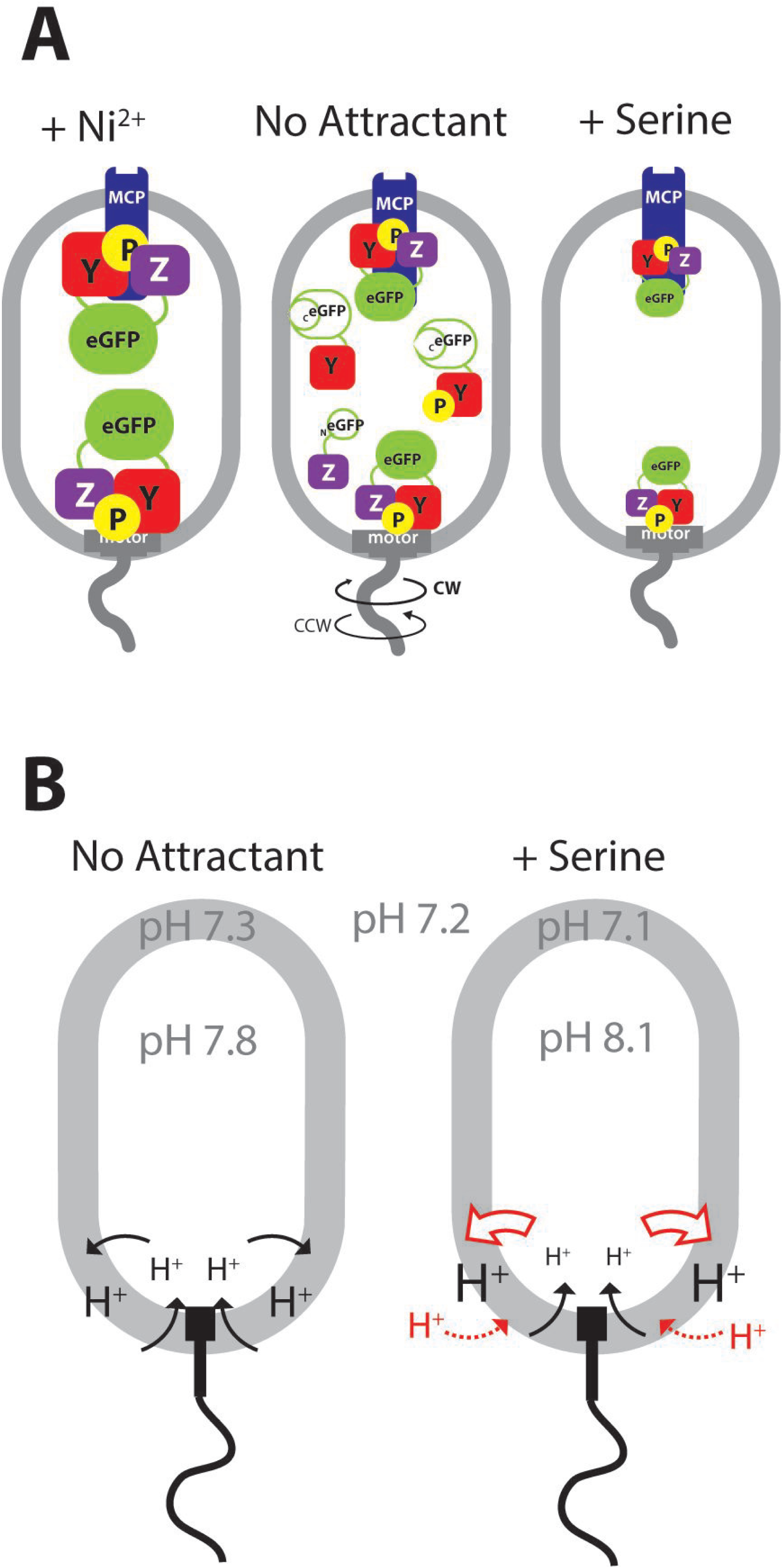
Model of observed split-eGFP and pHluorin behaviour in chemotactically-active *E. coli* cells. **(A)** Split-eGFP foci reconstituted from interacting CheY~P-_C_eGFP and CheZ-_N_eGFP both at chemoreceptor (MCP) as well as motor complexes. In cells exposed to Ni^2+^, there is a tendency for more and brighter eGFP foci. Cells perceiving serine tend to form less bright and fewer eGFP foci. For simplicity, non-phosphorylated CheY is not drawn in the schematic case of serine and Ni^2^+. **(B)** In absence of attractant, motile *E. coli* cells maintain a difference of ~0.8 pH unit between cyto- and periplasm. Cells accumulating in a radial gradient from a 100−μM serine source after 20 min increase cytoplasmic pH (pH 8.4), possibly to sustain higher proton flux through the flagellar motors.

## MATERIALS AND METHODS

**Bacterial strains and culture conditions**. A specific motile strain of *E. coli* MG1655, obtained from the *E. coli* Genetic Stock Center (Yale University; CGSC#8237), was used as host strain for the plasmids constructed in this study. For routine growth, *E. coli* was cultured on LB medium supplemented with the appropriate antibiotics, if necessary. For chemotaxis assays, the strains were grown at 37°C with 180 rpm shaking in M9-Glc-medium. M9-Glc consists of 17.1 g l^−1^ Na_2_HPO_4_-12H_2_O, 3.0 g l^−1^ KH_2_PO_4_, 0.5 g l^−1^ NaCl, 1.0 g l^−1^ NH_4_Cl, 4 g l^−1^ of glucose, 1 g l^−1^ of Bacto™ casamino acids (BD Difco), Hutner’s trace metals (54), and 1 mM MgSO_4_. The medium was supplemented with 30 μg chloramphenicol (Cm) ml^−1^ for strains containing pSTV-based plasmids and 100 μg ampicillin (Amp) ml^−1^ for plasmids expressing pHluorin. All used strains are detailed in Table 1.

**Cloning of the split-eGFP**. The chemotaxis regulator protein CheY was fused (at its C-terminal end) with the C-terminal part of eGFP, and its phosphatase CheZ was fused (at its C-terminal end) with the complementary N-terminal part of eGFP. First, the gene coding for *cheY* (without its stop codon) was amplified by PCR from *E. coli* MG1655 genomic DNA using primers 130719 and 130720, elongated with BamHI and AatI restriction sites, respectively (Table S1). The 3’-end of *egfp* (*_C_egfp*) corresponding to amino acids 158–238 was amplified by PCR from plasmid pPROBE (38) using primer 130724 containing an AatI restriction site and a sequence for a seven-amino-acid linker (GTSGGSG), and primer 130725, elongated with SpeI and HindIII restriction sites (Table 2). The *cheY* fragment was digested with BamHI and AatI, and the *_C_egfp* fragment with AatI and SpeI, and both were ligated downstream of the synthetic promoter P_AA_ in pSTV28P_AA_mcs, cut with BamHI and SpeI (55).

**Table 2.**
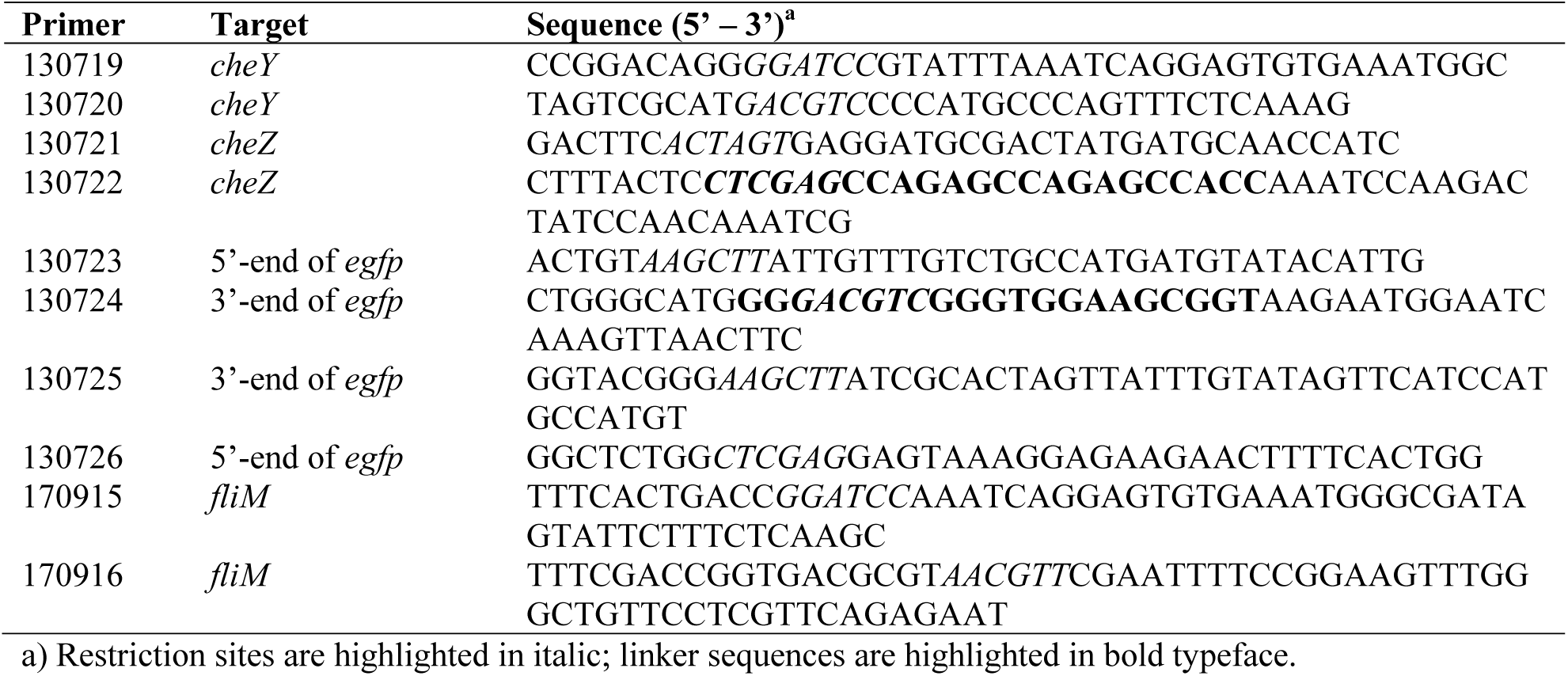
Primer list

In a second cloning step *cheZ* (without stop codon) was amplified by PCR from *E. coli* MG1655 genomic DNA using primer 130721 elongated with an SpeI site, and primer 130722 containing an XhoI restriction site and a sequence encoding an eight-amino-acid linker (GGSGSGSR). The 5’-end of *egfp* (*_N_egfp*) encoding amino acids 2−157 was amplified from plasmid pPROBE using primers 130726 and 130723, elongated with XhoI and HindIII restriction sites, respectively. Both PCR fragments were then inserted downstream of the *cheY-_C_egfp* hybrid gene within the same operon (i.e, under P_AA_ control), by digestion with SpeI and HindIII, and ligation. A variety of derivative plasmids was created, in which P_AA_ was replaced by different weaker synthetic promoters named P_JJ_, P_II_, and P_OO_ in order to tune the level of expression of the fusion proteins (42). Synthetized promoter sequences were flanked by EcoRI and BamHI restriction sites, to allow easy exchange (56). Equivalent constructs of the *P_JJ_-cheY-_C_egfp-cheZ-_N_egfp* were produced, in which LVA, AAV and ASV-instability tags were added to the 3’-end of *_C_egfp* (38) (Table 1). As negative controls we introduced frameshift mutations in *cheY* and *cheZ*, by digestion with the restriction enzymes SalI and MluI, respectively, filling in and religating.

To verify colocalization of reconstituted CheY-CheZ-eGFP foci with the flagellar motor, a *fliM-mCherry* translational fusion construct was produced as follows. The *fliM-*gene of *E. coli* was amplified by PCR using primers 170915 and 170916, appended with internal BamHI and MluI-sites, respectively. After PCR, the amplicon was digested with BamHI and MluI, and ligated with plasmid pJJUN-L-mche (56), cut with the same enzymes. This links the *fliM* coding region via a short linker to the *mCherry* coding frame. After transformation into *E. coli* and verification by sequencing this resulted in plasmid p5920. The plasmid was purified and cotransformed into *E. coli* 4703 (Table 1).

*E. coli* deletion mutants were constructed by double recombination methods (57). Gene flanking regions were amplified by PCR and cloned into the pEMG suicide vector (57) and transformed into *E. coli* MG1655. Single recombinants were recovered by selection on Km-resistance and were examined by PCR. If correct, they were transformed with the plasmid pSW-I that contains the gene for the ISceI restriction enzyme (57). Induction of ISceI expression by 3-methylbenzoate forces the second recombination. Kanamycin-sensitive colonies were examined by PCR for proper deletion. Finally, pSW-I was cured by consecutive passage in culture without antibiotic selection pressure. Notably, we deleted *fliC*, *fliM, cheA, tsr* and *cheY-cheZ* (Table 1).

**Construction of pHluorin fusions**. In order to express pHluorin in the periplasm, the *torA* export signal sequence (58) was added to the 5’-end of the *pHluorin* sequence (AF058694.2) flanked by two NcoI restriction sites. The DNA sequence was synthetized by DNA 2.0 (CA, USA) and delivered in their custom vector pJ201. The complete synthetized gene sequence was flanked with EcoRI and HindIII restriction sites. The *torA-pHluorin* sequence was cloned downstream of P_BAD_ in pBAD24 (59) using EcoRI and HindIII restriction sites to express the protein after induction with arabinose. After transformation in *E. coli* MG1655 this yielded plasmid pCRO4. From pCRO4, the *torA* signal sequence was removed by digestion with NcoI, after which the plasmid was religated. This resulted in plasmid pCRO5, which was used to express pHluorin in the *E. coli* cytoplasm.

**Epifluorescence microscopy of reconstituted split-eGFP fusions**. Overnight cultures of *E. coli* strains in M9-Glc supplemented with 30 μg Cm ml^−1^ were 100–times diluted in fresh medium and incubated at 37°C with shaking until they reached an optical density at 600 nm (OD_600_) of 0.5–0.7. From these exponential cultures, an aliquot of 600 μl was centrifuged at 2,400 × g for 5 min to collect the cells (note that motile *E. coli* cells do not form a strong pellet). A volume of 500 μl of supernatant was carefully removed and 1 ml of motility buffer was added to the remaining 100 μl cell suspension (motility buffer is 10 mM potassium phosphate, 0.1 mM EDTA, 10 mM lactate, 1 μM methionine, pH 7.0) (60). This procedure was repeated once and finally the cells were concentrated in 50 μl of remaining motility buffer. A drop of 7 μl of cell suspension was spotted on 1% agarose (in motility buffer) 1 mm–thick coated microscopy slides and then covered with a regular 0.17–mm thick glass coverslip. Cells were imaged at an exposure time of 10 ms (phase-contrast) or 1 s (eGFP) with a DFC 350 FX R2 Leica camera mounted on an inverted DMI 4000 Leica microscope using a 100× Plan Apochromat oil objective. The images were analyzed using the open-access software *MicrobeJ*, which allows automatic cell segmentation and foci detection (www.microbeJ.com) (61).

**Time-lapse microscopy**. Dynamic eGFP foci behaviour was followed by time-lapse microscopy. Overnight cultures of *E. coli* strain 5395 (Δ*cheYcheZ* + pCRO9) or 5430 (Δ*cheYcheZ* + pCRO32) in M9-Glc supplemented with 30 μg Cm ml^−1^ were 100–times diluted in fresh medium, and incubated at 37°C with shaking until they reached an OD_600_ of 0.5–0.7. From these exponential cultures, an aliquot of 1 ml was centrifuged at 2,400 × g for 5 min to concentrate the cells. The concentrated cells were washed twice with 1 ml of motility buffer as above, and finally concentrated in 600 μl of motility buffer. Cells were allowed to adhere to the bottom of the wells of a CELLview™ Slide with 10 wells (Greiner Bio-One GmbH, Germany) coated with poly-L-lysine. For coating, 100 μl of 0.01 % poly-L-lysine solution (Sigma-Aldrich) was incubated for 5 min at room temperature in the wells, decanted, after which the wells were dried overnight. Wells were filled with 100 μl of cell suspension (see above) and incubated at room temperature for 30 min. Liquid was decanted and wells were washed 5 times with 400 μl of motility buffer to remove any non-adhering cells. Finally, the wells were filled with 200 μl of motility buffer. Cells were imaged with a Nikon Eclipse Ti-E inverted microscope, equipped with an ORCA-flash4.0 camera (Hamamatsu) and a Plan Apo λ 100×1.45 oil objective (Nikon). Seven regions of the wells were imaged automatically with one minute intervals for eGFP fluorescence (200 ms exposure) and phase contrast (10 ms exposure) with lamp power at 100 % (Lumencor). Baseline eGFP fluorescence was imaged at time=0 and 1 min, after which 10 μl of inducer (2 mM of NiCl_2_ or serine to a final concentration of 100 μM, or motility buffer as control) was pipetted in the wells, and cells were imaged for a further 10 min (at 1 min intervals). Images were recorded using Micro-Manager 1.4 (http://www.micro-manager.org/).

**Time-lapse data analysis**. Ellipses of 8×8 pixels were placed on visible foci using MetaMorph (Series 7.5, MDS, Analytical Technologies) and mean fluorescence intensities in the ellipses were extracted at all time points using an in-house written Matlab (R2015b) script (developed by Serge Pelet, University of Lausanne). Baseline fluorescence decay of foci in individual cells incubated with motility buffer only was fitted with a general second-order decay function using Igor Pro (WaveMetrics, Inc. Oregon 97035, USA), as suggested in Ref (62):

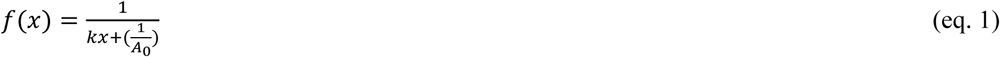
with *k* being the rate constant and *A_0_* the initial fluorescence. The average rate constant (AVE_k,buffer_) and its standard deviation (SD_k,buffer_) were calculated from all individual cell traces in motility buffer. Individual fluorescence responses of the foci of experiments with induction of nickel or serine were fitted with the same decay function (between time points 2 – 12 min), obtaining *k_Ni_* or *k_Ser_* and their fitting deviations SD*_k,Ni_* or SD*_k,Ser_*.

The number of deviant individual response curves was then counted for each condition, motility buffer only, nickel or serine, if *k_Ni or Ser_ + SD_k_,_Ni or Ser_ < AVE_k_,_buffer_ − SD_k,buffer_*, which corresponds to a fitting curve that decreases less than the expected baseline second-order decay or if *k_Ni or Ser_ + SD_k,Ni or Ser_ > AVE_k,buffer_ – SD_k,buffer_*, which corresponds to a fitting curve that decreases more than the expected baseline second-order decay. The proportions of individual deviant curves in a simultaneous sampling series between buffer incubation and nickel or serine induction were then compared using a Kruskal-Wallis test.

Foci intensities and positions of reconstituted split-eGFP and FliM-mCherry were further automatically extracted from cell images using SuperSegger (63), which subtracts local background fluorescence from the cell, and summarized by custom-written Matlab scripts to calculate the foci localization and intensity distributions. A focus score of 10 was used as threshold for foci calculations. Changes in foci distributions over time of cell exposure to attractant or repellent were tested on linear regression of slopes for the respective bin-size, for treatment against motility buffer only (Fig. S1, S2), as recommended by Ref (64).

**Expression of pHluorin for epifluorescence microscopy assays**. Overnight cultures in Mg-Glc Amp medium of *E. coli* MG1655 (pCRO5) or MG1655 (pCRO4) were diluted in the same medium supplemented without or with a range of L-arabinose concentrations (1%, 0.1%, 0.01%, 0.001% *w/v*), and were further incubated at 37°C until reaching exponential phase (OD_600_ ~ 0.5). The *E. coli* MG1655 (pCRO5) culture was then sampled directly for epifluorescence microscopy of pHluorin expression. To allow export of TorA-pHluorin into the periplasm, strain MG1655 (pCRO4) was harvested in exponential phase by centrifugation at 2,400 × g for 5 min to pellet the cells. Cells were resuspended in the same volume of M9-Glc without arabinose and incubated for a further 2 h at 37°C, after which they were sampled for observation of TorA-pHluorin expression.

Culture aliquots of 400 μl culture were centrifuged at 2,400 × g for 5 min to pellet the cells, which was resuspended in 50 μl of M9-Glc medium without arabinose. A drop of 7 μl of this cell suspension was deposited on a 1% agarose in M9 medium coated standard microscopy slide and covered with a cover slip. Images were immediately taken with a Zeiss Axioplan II epifluorescence microscope equipped with a 100× Plan Apochromat oil objective (Carl Zeiss, Jena, Germany) and a SOLA SE light engine (Lumencor, USA). We used an eGFP HQ excitation filter (470 nm, 40 nm bandwidth) and an emission filter at 525 nm (bandwidth 50 nm, Chroma Technology Corp., VT, USA).

To test dependency of pHluorin fluorescence on external pH, cultures were grown as above, centrifuged, but resuspended in 50 μl of M63 minimal medium supplemented with 2 g l^−1^ casein hydrolase, 20 mM sodium benzoate and buffer to the respective test pH. M63 medium consists of 0.4 g l^−1^ KH_2_PO_4_, 0.4 g l^−1^ K_2_HPO_4_, 2 g l^−1^ (NH_4_)_2_SO_4_ and 7.45 g l^−1^ KCl. To obtain pH 6.0, we used 50 mM 2-(N-morpholino)ethanesulfonic acid (MES); for pH 7.0, we used 50 mM (3-(N-morpholino)propanesulfonic acid (MOPS); and for pH 8.0, we used 50 mM of N-Tris(hydroxymethyl)methyl-3-amino-propanesulfonic acid (TAPSO) (65). A drop of 7 μl of bacterial suspension was then deposited on a standard microscopy glass slide coated with a 1% agarose solution in the corresponding M63-medium and pH. Cells were immediately imaged in phase-contrast (10 ms) and epifluorescence (100 ms, below) using an sCMOS camera (Flash4.0, Hamamatsu) mounted on an inverted Ti-Eclipse epifluorescence microscope (Nikon) using an Apo PLAN 100×1.45 oil objective. The excitation was provided by a solid-state light source (SpectraX, Lumencor) at a wavelength of 386 nm (bandwidth: 23 nm) or at a wavelength of 470 nm (bandwidth: 40 nm), and fluorescence emission was detected at a wavelength of 525 nm (bandwidth: 30 nm).

**Source attraction assays**. Active bacterial chemotactic response was obtained and deduced from modified agarose source attraction assays, adapted for microscopy observations. Briefly, on a microscopy glass slide (1 mm thickness, RS France, Milian), two squared coverslips (24 × 24 mm, 0.17−mm thickness) were deposited on the edge of the slide. A 5 μl drop of 2% agarose supplemented with 100 μM of serine (kept at 55°C) was pipetted on the slide between the coverslips. A large coverslip (25 × 50 mm, Menzel-Glaser) was quickly deposited on top of the agarose plug. After drying during 5 min, 150 μl of washed exponentially-growing *E. coli* cell suspension in motility buffer were pipetted at the edge between the large coverslip and the glass slide. Cell accumulation was measured after 30 min by light microscopy close to the agarose plug source as outlined elsewhere (66).

Cells expressing pHluorin were imaged close and on either side of the agarose source after 15 to 35 min incubation at room temperature, using an sCMOS camera (Flash4.0, Hamamatsu) mounted on an inverted Ti-Eclipse epifluorescence microscope (Nikon) equipped with an N PLAN 20× air objective and wavelength settings as described above. Exposure times for pHluorin in this case were 30 ms for both fluorescence channels. Four independent source or negative control replicates were produced for every strain. Cells were identified and their abundance was quantified using the “find edges” routine in ImageJ. The intensity values were averaged across successive outward-moving sectors of 25 pixels width (corresponding to 2 μm) parallel to the border of the agarose source (3 zones within the source and 57 zones outside the plug). Fluorescence values were extracted using the same sectors on the respective images. Intensity values were then averaged on both sides and across four replicates, and plotted as a function of distance ± one *SD*.

A modified agarose plug assay was developed to measure dynamic split-eGFP reconstituted foci intensity distributions in individual cells exposed to attractant or repellent gradient. A 1–mm thick flexible glass slide support was used to form an ellipsoid of approximately 4 cm in length, consisting of 1% agarose in motility buffer. After solidifying, one third of this was cut out and replaced by a 1% agarose source in motility buffer alone, or containing 100 μΜ serine or 100 μΜ NiCl_2_. After solidifying for 12 min, exponentially growing, washed and resuspended *E. coli* strain 5430 cells (Δ*cheYcheZ* pCRO32) in motility buffer were spread on the agarose surface next to the source area. Cells were imaged for eGFP foci at between 10 and 30 min after application, at three X-positions at relative distances of 8 mm from each other, and five Y-positions 300 μm apart.

## ACKNOWLEDGEMENTS

The authors thank Lavinia Bottinelli and Giulia Torriani for their help in the early phases of split-eGFP and pHluorin cloning experiments. This work was supported by the Swiss National Science Foundation NanoTera project 20NA21-501 143082, and by financing from the Herbette Foundation (2018-1-D-26). The authors declare no conflict of interest.

